# Mechanism of auxin and GA crosstalk in diploid strawberry fruit initiation

**DOI:** 10.1101/2020.03.30.016188

**Authors:** Junhui Zhou, John Sittmann, Lei Guo, Xiaolong Huang, Anuhya Pulapaka, Zhongchi Liu

## Abstract

Strawberry, a high value fruit crop, has recently become amenable for genetic studies due to genomic resources and CRISPR/CAS9 tools. Unlike ovary-derived botanical fruits, strawberry is an accessory fruit derived from receptacle, the stem tip subtending floral organs. Although both botanical and accessory fruits initiate development in response to auxin and GA released from seeds, the downstream auxin and GA signaling mechanisms underlying accessory fruit development remain unknown. Using wild strawberry, we performed in depth molecular characterizations of accessory fruit development. We show that auxin signaling proteins FveARF8/FveARF6 are bound and hence inhibited by FveIAA4 and FveRGA1, repressors in auxin and GA signaling pathways. This inhibition is relieved post-fertilization or by the application of GA or auxin. Mutants of *FveRGA1* developed parthenocarpic fruit suggesting a conserved function of DELLA proteins in fruit set. Further, FveARF8 was found to repress the expression of a GA receptor gene GID1c to control fruit’s sensitivity to GA, revealing a novel crosstalk mechanism. We demonstrate that consensus co-expression network provides a powerful tool for non-model species in the selection of interacting genes for functional studies. These findings will facilitate the improvement of strawberry fruit productivity and quality by guiding future production of parthenocarpic fruit.

**One sentence summary:** Genome editing and network analysis are applied to investigating the mechanism of accessory fruit initiation in the wild strawberry, revealing conserved as well as novel crosstalk mechanisms.

The author responsible for distribution of materials integral to the findings presented in this article in accordance with the policy described in the Instructions for Authors (www.plantcell.org) is: Zhongchi Liu

## Introduction

Strawberry is a high-value fruit crop immensely important for agriculture and human nutrition. Unlike most fruits developed from the ovary wall, strawberry fruit is derived from receptacle, the stem tip supporting floral organs (Hollender et al., 2012). About 200 achenes, ovaries each with a single seed inside, dot the surface of the receptacle, making them highly accessible for manipulation. Nitsch’s 1950 experiment showed that removal of achenes from a strawberry receptacle prevented receptacle fruit enlargement. However, exogenous application of auxin could substitute for the achenes and stimulated receptacle fruit growth, indicating achenes as the source of auxin (Nitsch 1950). Later work showed that emasculated wild strawberry flowers could still develop into fruit if auxin or GA, alone or in combination, was supplied exogenously (Kang et al., 2013; Liao et al., 2018). Transcriptome analyses and reporter gene expression confirm that fertilization-induced biosynthesis of auxin and GA occurs in the embryo and endosperm (Kang et al., 2013; Feng et al., 2019). The phytohormones are then transported to the receptacle to stimulate fruit enlargement. Exactly how auxin and GA interact to coordinately regulate receptacle fruit development is not known.

The fertilization-induced phytohormone synthesis for “fruit set”, the no return commitment of the floral tissue to enter fruit pathway, is an evolutionarily conserved strategy in angiosperms. The unique strawberry fruit structure as well as prior research provide a strong foundation for further investigations into signal communication between achenes and the receptacle and auxin/GA crosstalk during fruit development. This is further aided by the rich genomic resources including genome sequencing of wild strawberry *Fragaria vesca* and garden strawberry *Fragaria x ananassa* (Shulaev et al., 2011; Edger et al., 2018; Edger et al., 2019) as well as extensive transcriptome data and network analyses during flower and fruit development (Kang et al., 2013; Hollender et al., 2014; Sanchez-Sevilla et al., 2017; Shahan et al., 2018). As garden strawberry is an allo-octoploid, wild strawberry *F. vesca,* a diploid species, is used in this study.

Prior research in model organisms illuminated auxin and GA signaling cascade. When auxin is absent, IAAs bind to ARFs to prevent ARFs from activating or repressing downstream target genes. When auxin is synthesized or supplied, it binds its receptors TIR/AFBs, which are F-box proteins that recruit and add ubiquitins to the Aux/IAA proteins. This leads to the 26S proteasome-mediated degradation of Aux/IAAs and the release of ARFs to execute their functions (Leyser, 2018). Similarly, in the absence of GA, DELLA repressor proteins accumulate and bind transcription factors to prevent their function. When GA is produced, GA binds to its receptor GID1 to facilitate binding of GID1 to DELLA that leads to ubiquitination and proteasome-mediated degradation of DELLA. Transcription factors previously bound by DELLA are now free to act upon downstream genes (Daviere and Achard, 2013).

In model systems Arabidopsis and tomato (both with botanical fruits), fertilization-independent fruit formation (parthenocarpy) could be induced when repressors of auxin or GA signaling response are mutated or reduced as demonstrated by *arf8* mutants in *Arabidopsis* and transgenic knockdowns of tomato *ARF7, ARF8,* or *IAA9* (Wang et al., 2005; Goetz et al., 2006; Goetz et al., 2007; de Jong et al., 2009; Wang et al., 2009). These results led to the proposal that the IAA9/ARF7/ARF8 repressor complex inhibits fruit set in the ovary tissue of Arabidopsis and tomato. To investigate the relationship between auxin and GA, several studies were carried out in Arabidopsis, pea, and tomato which showed that auxin acts upstream of GA by activating the biosynthesis of GA (Serrani et al., 2008; Dorcey et al., 2009; Ozga et al., 2009; Fuentes et al., 2012). Recently, direct protein-protein interactions between SlDELLA and SlARF7 and SlIAA9 were shown to serve as another mechanism mediating crosstalk between auxin and GA in tomato fruit (Hu et al., 2018).

In contrast, there has been little functional studies of fruit set in accessory fruits including strawberry. Previously, we employed RNA-seq to profile finely dissected strawberry fruit tissues, embryo, ghost (seed coat + endosperm), ovary wall, and receptacle pith and receptacle cortex, at five early fruit developmental stages (Kang et al., 2013). Based on the RNA-seq data and subsequent reporter GUS studies (Feng et al., 2019), the endosperm within the achene emerges as the site of fertilization-induced auxin and GA biosynthesis targeted for fruit development. However, auxin and GA perception and signaling genes, TIR/IAAs and GID1c/RGA1, are highly expressed in the receptacle, indicating receptacle as the site of auxin and GA perception and response (Kang et al., 2013). Using the RNA-seq data described above, consensus co-expression network was established (Shahan et al., 2018), which identifies robust co-expression modules in receptacle fruit development. In addition, CRISPR/Cas9 genome editing has been demonstrated in wild and garden strawberries (Zhou et al., 2018; Feng et al., 2019; Martin-Pizarro et al., 2019), providing tools for dissecting gene function.

Here, we provide an in depth functional dissection of a few key auxin and GA signaling components in the receptacle fruit development in diploid strawberry. We observed both conserved and novel cross-talk mechanisms for GA and auxin signaling, identified *FveRGA1* as a key repressor of fruit set in strawberry, and validated the usefulness of consensus co-expression network. The findings lay the foundation for future improvement of strawberry fruit.

## Results

### *arf8* mutants develop larger and rounder fruits

Previously, we reported successful genome editing of *FveARF8,* and the *arf8* mutants showed faster seedling growth (Zhou et al., 2018). However, *arf8* adult plants are similar to WT plants in stature. To determine the defect of *arf8* mutants in flower and fruit development, we first characterized four independent CRISPR lines of *arf8,* two lines are *arf8* (−2, −2), one line is *arf8* (−2+1, −2+1), and one line is *arf8* (−8, −23+5) (Figure S1). Second, we established that fruits derived from primary and secondary flowers are similar in size (Figure S2, allowing us to collect data from both primary and secondary fruits. Third, to remove impact of environment on plant vigor and hence fruit size, the height and width of fruits were normalized respectively to the height and width of petals of the same flower.

*arf8* mutant fruit is larger and rounder than wild type (Figure 1). Significant increase in the width and the height of *arf8-1 (−2, −2)* fruit is found when compared with wild type (Figure 1A, B); the higher increase of fruit width results in rounder shape in developing fruit. The other three CRISPR-induced *arf8* lines exhibited similar phenotype as *arf8-1* with increased fruit height and width (Figure 1C). Hence, *arf8-1* is used for the subsequent analyses. In ripe fruit, the width remains larger in *arf8-1* mutants, but the height becomes similar to WT (Figure 1A, D) probably due to compensatory mechanisms at late stages of fruit development. *arf8* mutants also exhibited a mild floral phenotype; they showed an increase in petal number (Figure S2C). While 80% wild type flowers have 5 petals, only 40% *arf8* mutants produce 5 petals and the remaining flowers produce 6 or 7 petals (Figure S2D). The average petal size however appears the same between WT and *arf8-1* mutants (Figure S2E).

**Figure 1.**
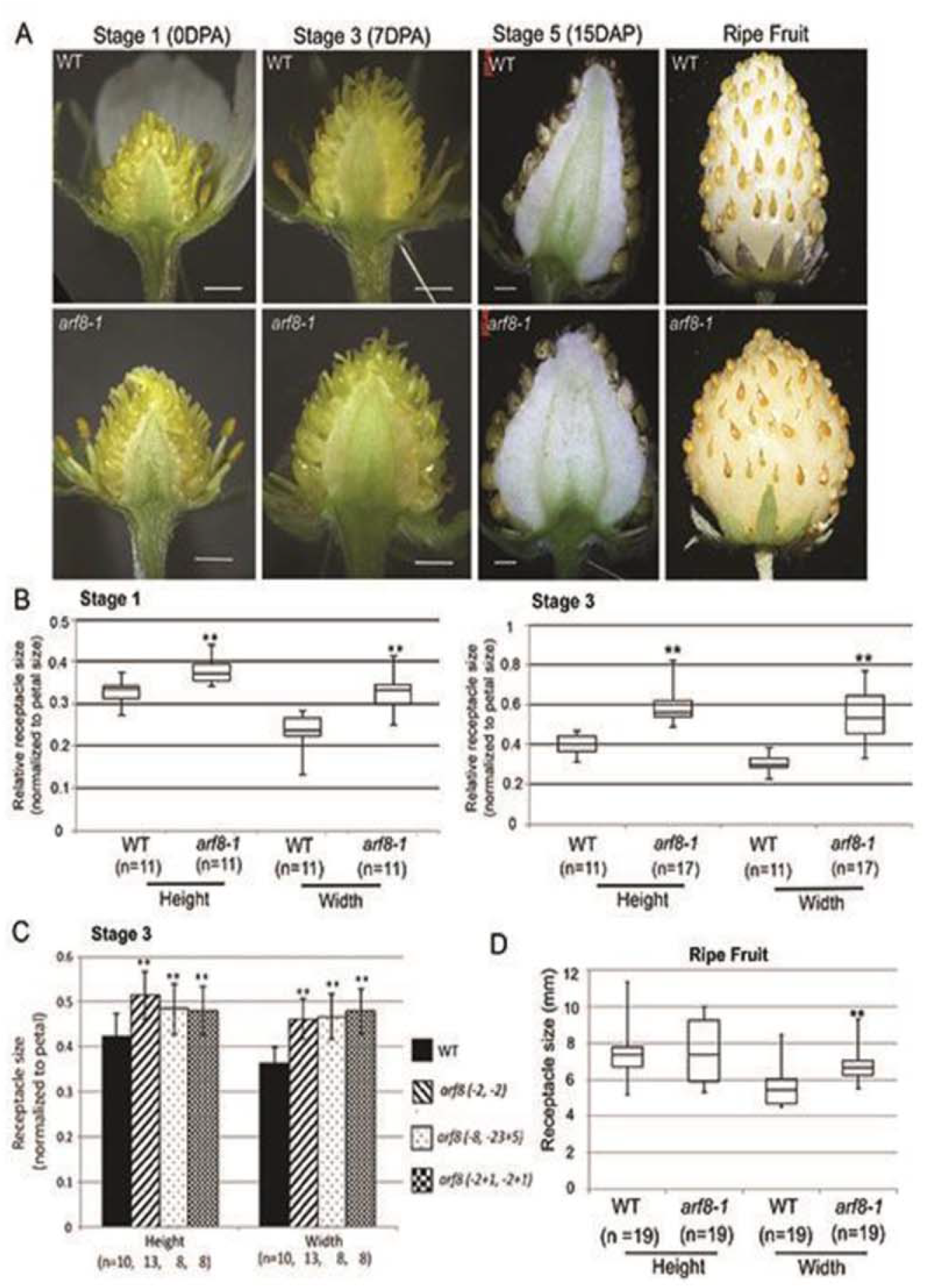
*arf8* mutants develop larger and rounder fruit. A. Images of bisected receptacle at stage 1 (0DPA, before fertilization), stage 3 (7DPA, post-fertilization), stage 5 (15 DPA), and ripe stage in WT and *arf8-1 (−2, −2).* Note the rounder fruit shape in *arf8-1.* Bars; 1 mm. B. Average size of receptacle fruit (height and width, normalized to petal size) at stage 1 and stage 3. ** above bars represent significant difference (P < 0.01) between WT and *arf8-1* mutant. C. Relative receptacle fruit size (height and width, normalized to petal size) in WT and three additional *arf8* mutant lines at stage 3 (7DPA). These *arf8* mutant lines, #156 (−2, −2), #391-3 (−8, −23+5), and #391-31 (−2+1, −2+1) are illustrated in Figure S1. D. WT and *arf8* mutant receptacle fruits at ripe stage. Since petal has senesced, the absolute fruit size was measured. *arf8-1* mutants show a significant increase in width but not in height at the ripe stage as shown in A.

Using the *Arabidopsis* Ubiquitin 10 promoter, *FveARF8* was over-expressed in *F. vesca*; the resulting transgenic lines (#1, #5, and #26), which exhibit a significantly higher *FveARF8* transcript level than the non-transgenic control, showed retarded seedling and plant growth (Figure S3). Transgenic lines #1 and #26 showed a reduction of fruit size when normalized to petal (Figure S3C, D), while transgenic line #4 with a wild type expression level of *FveARF8* exhibited normal plant statue (Figure S3A, B). Together, our data suggest that *FveARF8* is a negative regulator of plant growth and fruit enlargement.

### Mechanism of auxin and GA signaling in fruit set

Fertilization-induced phytohormone synthesis and signaling are critical for “fruit set”, a decision to proceed with fruit development upon successful fertilization (Gillaspy et al., 1993). In *F. vesca,* we previously showed that auxin and GA each substituted for fertilization to stimulate fruit set, yet each hormone alone stimulated fruit development to a size smaller than the pollinated control fruit (Kang et al., 2013). Only when both auxin and GA were applied at the same time, did parthenocarpic strawberry develop to the same size as the pollinated control, suggesting the requirement of both auxin and GA for strawberry fruit set. Nevertheless, the molecular mechanism underlying this double requirement is not known.

To test if *FveARF8* could act to block fruit set pre-fertilization, we examined *arf8-1* fruit formation in the absence of fertilization. Wild type and *arf8-1* mutant flowers were emasculated to prevent fertilization. Fruit size at 0DPA (pre-fertilization) was compared to fruit size at 7 DPA (post-emasculation) within the same genotype. No significant change in fruit size was observed at 7DPA for *arf8-1* as well as WT (Figure 2), suggesting that fertilization-induced hormone production is still required for *arf8* mutants to develop fruit and that *FveARF8* is unlikely involved in preventing fruit set.

**Figure 2.**
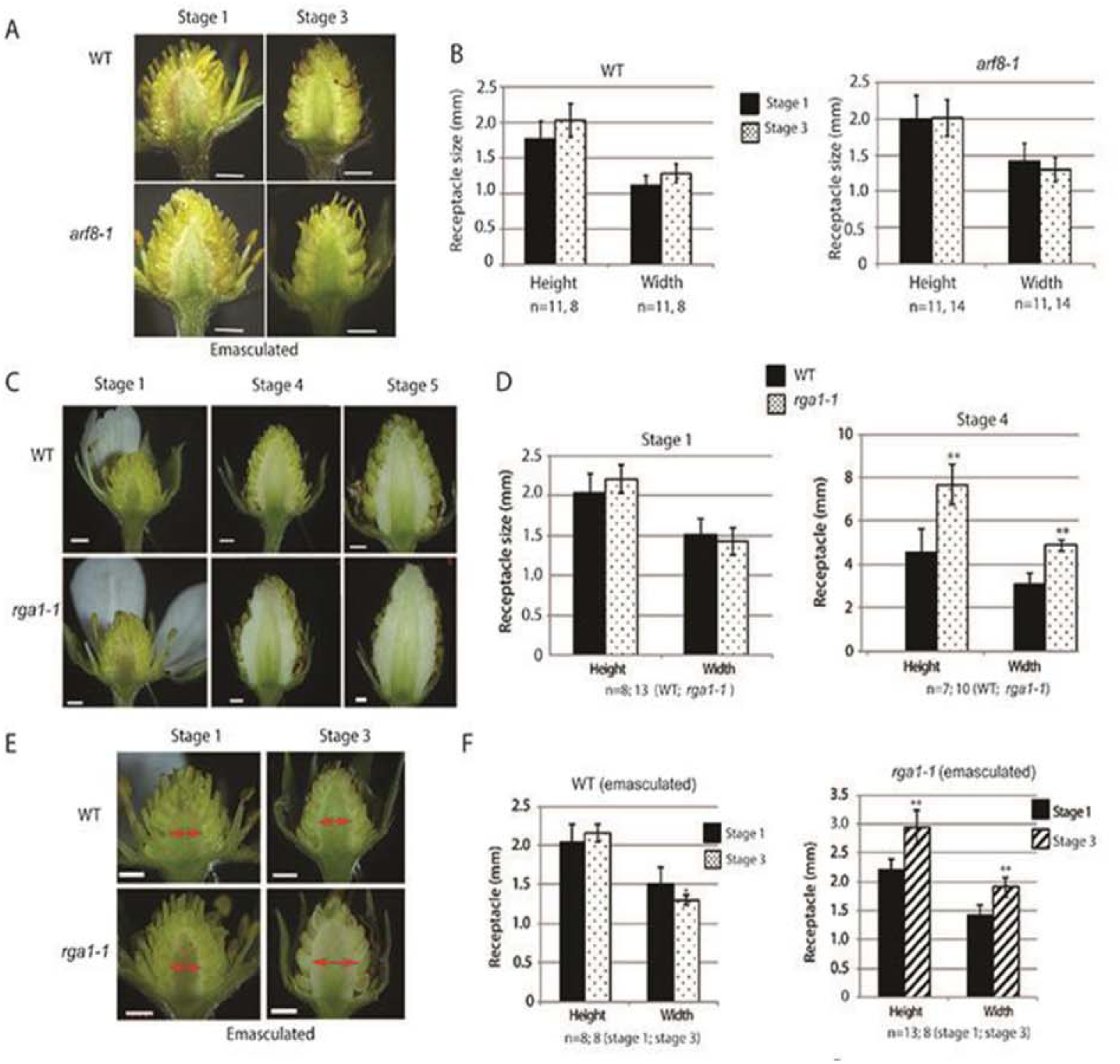
*rga1-1, but not arf8-1,* develop parthenocarpic fruit. A. Photos of bisected receptacle of WT and *arf8-1* at 0DPA and 7DPA. The flowers were emasculated at the 0DPA. Bars; 1 mm B. Quantitative measurement of receptacle size in emasculated fruit shown in A. No significant difference in fruit size between 0DPA and 7DPA was observed in emasculated WT and *arf8* receptacle. C. Images of bisected WT and *rga1-1* receptacle at stage 1 (1DPA), Stage 4 (9DPA), and stage 5 (11-12 DPA) without emasculation. *rga1-1* receptacle fruit is larger and longer than WT at stages 4-5. Bar: 1 mm. D. Quantitative measurement of WT and *rga1-1* receptacles at stage 1 and stage 4. *rga1-1* develops significantly larger fruit, particularly in receptacle height. E. Image of emasculated WT and *rga1-1* receptacles at stage 1 (1DPA) and stage 3 (7DPA). Red arrows highlight the width of the receptacle. Fertilization-independent fruit enlargement is evident in *rga1-1* mutant. Bar, 1 mm. F. Quantitative measurements of emasculated WT and *rga1-1* receptacles at stage 1 (0DPA) and stage 3 (7DPA). Emasculated WT receptacles at stage 1 and stage 3 are similar in size. Emasculated *rga1-1* receptacles are significantly larger and taller at stage 3 when compared with stage 1.

During GA signaling, the GA-GID1 receptor complex targets the DELLA repressor protein for proteasome-mediated degradation, resulting in the release of downstream transcription factors (TFs) normally sequestered by the DELLA protein (Hartweck, 2008; Sun, 2011). In *F. vesca, gene06210 (FveRGA1)* encodes one of the five annotated probable DELLA proteins and is the only one with a full DELLA motif (Caruana et al., 2018). A loss-of-function mutation *rga1-1* (also named *srl-1)* was identified in *FveRGA1* during a chemical mutagenesis screen (Caruana et al., 2018) (Figure S1B). This mutant offers an opportunity to determine if the DELLA repressor encoded by *FveRGA1* regulates fruit set in strawberry. *rga1-1* receptacle size was measured at Stage 1 (1DPA), stage 4 (9DPA), and stage 5 (11-12 DPA). While the receptacle size difference between WT and *rga1-1* is not obvious at 1DPA, the receptacle becomes significantly larger and taller at post-fertilization stages 4-5 (9DPA and 11-12 DPA) (Figure 2C, D), suggesting that constitutive GA response in the *rga1-1* receptacle may underlie the larger fruit phenotype. To determine if *rga1-1* mutant could develop parthenocarpic fruit, emasculated WT and *rga1-1* flowers were marked and the receptacle size measured. While WT fruit remains unchanged when measured at stage 1 (0DPA) and stage 3 (7DPA), *rga1-1* mutants develop significantly larger receptacle at stage 3 (7DPA) when compared with stage 1 (0DPA) (Figure 2E, F). The data suggests that *FveRGA1* encodes a key negative regulator of fruit set, perhaps by binding and inhibiting the activity of unknown TFs.

To investigate if the increased fruit size *in arf8* or *rga1-1* is due to increased cell division or cell expansion, histological sections were conducted. Histological sections of wild type, *arf8-1* and *rga1-1* fruits at 7DPA reveal an increased number of cell files across the width of the pith in *arf8* and *rga1-1* receptacle (Figure S4); WT (YW5AF7) has an average of 28 cell files, *arf8-1* has an average of 40 cell files, and *rga1-1* has an average of 46.5 cell files across the width of the pith (Figure S4J). In addition, *arf8-1* and *rga1-1* appear to have dense cell files near the vasculatures (Figure S4F, I). Thus, increased cell division might have contributed to the increased fruit width in both *arf8* and *rga1-1* mutants.

### *FveRGA1,* a key crosstalk node and regulator of fruit set

To investigate the mechanism of how auxin and GA coordinate their effect on fruit set and fruit growth and explain distinct effects of *arf8-1* and *rga1-1* on parthenocarpy, we investigated if FveRGA1 interacts with FveARF8 and FveARF6. FveARF6 was included as it was previously shown in *Arabidopsis* to interact with and inhibited by RGA1(Oh et al., 2014). In BiFC, FveRGA1 interacts positively with FveARF6 and FveARF8 (Figure 3). In Y2H, full length FveRGA1 self-activates, thus a N-terminal 564 bp deletion was made leading to FveRGA1-C (similar to the M5 version of AtRGA in Arabidopsis). Consistent with the BiFC result, FveRGA1-C interacts positively with FveARF6 and FveARF8 (Figure 3B), suggesting FveRGA1 as a key crosstalk node for hormone signaling.

**Figure 3.**
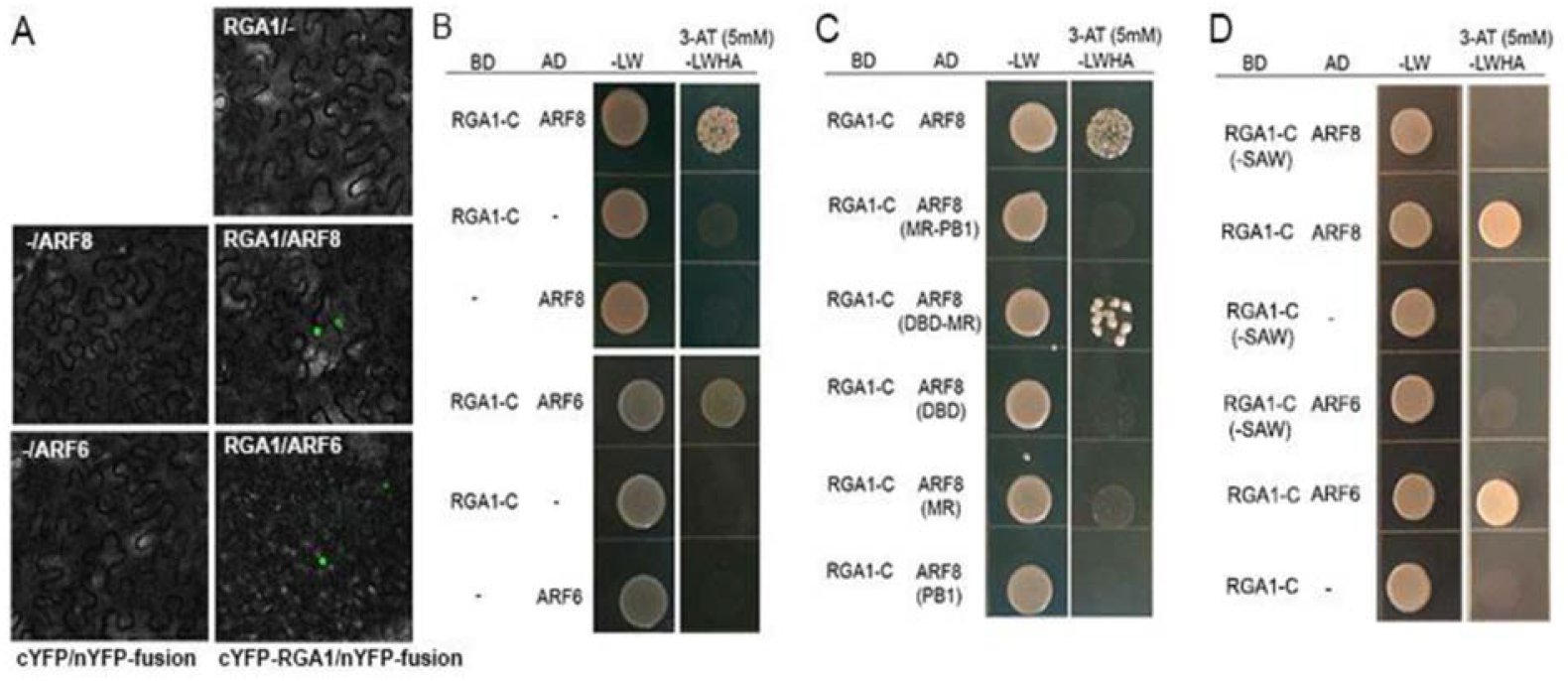
BIFC and Y2H revealed direct interaction of FveRGA1 with FveARF8, and FveARF6. A. BiFC in tobacco leaf showing positive interactions between FveRGA1 and respectively FveARF8 and FveARF6. B. Y2H assay confirming the interactions between FveRGA1-C (fused to BD) and respectively FveARF8 and FveARF6. Because of self-activating activity, a truncated FveRGA1-C with its N-terminal DELLA domain removed is used in the Y2H. C. Dissection of FveARF8 protein domains reveals the requirement of both DBD and MR for the interaction with RGA1-C. D. Removing the SAW domain from RGA1-C eliminates its ability to interact with FveARF6 and FveARF8 in Y2H assay.

To determine which domain of FveARF8 interacts with FveRGA1, FveARF8 is divided into three segments: DBD (DNA-binding), MR (Middle Region), and PB1 (Figure S5). While each domain of FveARF8 fails to interact with FveRGA1-C, the combined DBD-MR domains of FveARF8 positively interact with FveRGA1-C (Figure 3C). The work also indicates that the C-terminal portion of FveRGA1 is sufficient for the interaction with these TFs. Since *rga1-1* is a W601X mutation that deletes the last 10 amino acids from the C-terminal SAW domain (Figure S1B), we tested if removing the SAW domain in FveRGA1-C affect the interaction with FveARF8 or FveARF6 in Y2H. FveRGA1-C (-SAW) failed to interact with FveARF8 and FveARF6 (Figure 3D). *rga1-1* may thus fail to inhibit fruit set due to its inability to bind and inhibit FveARF6, ARF8, and perhaps other unidentified transcription factors.

The emerging theme from above studies is that *FveRGA1* acts to prevent fruit set (transition from stage 1 to stage 2). Fertilization induced GA production may remove FveRGA1 through GA-GID1-induced degradation, permitting fruit set to occur accompanied by the activation of a number of TFs.

### FveARF8 represses the expression of GA receptor *FveGID1c*

Since *arf8-1* fruit could not develop in the absence of fertilization (Figure 2A), the larger fruit of *arf8-1* post-fertilization (Figure 1A) may reflect an increased sensitivity or enhanced response to fertilization-induced hormonal signals, namely auxin and/or GA. To test this hypothesis, emasculated flowers of *arf8-1* were treated with 1mM NAA (1-naphthaleneacetic acid) or 1mM GA3. Mock treated fruits of the same genotype served as the control. The ratio of fruit size between hormone treated vs. mock-treated fruit at 7DPA was compared; the ratio between treated and mock for *arf8-1* mutants is significantly higher than that of WT under GA treatment as well as under auxin (NAA) treatment (Figure 4). Therefore, *arf8-1* mutant may exhibit higher sensitivity or response to GA and NAA than wild type, suggesting that one normal function of *FveARF8* is to dampen the receptacle’s sensitivity or response to GA and NAA perhaps in a feedback regulatory loop.

**Figure 4.**
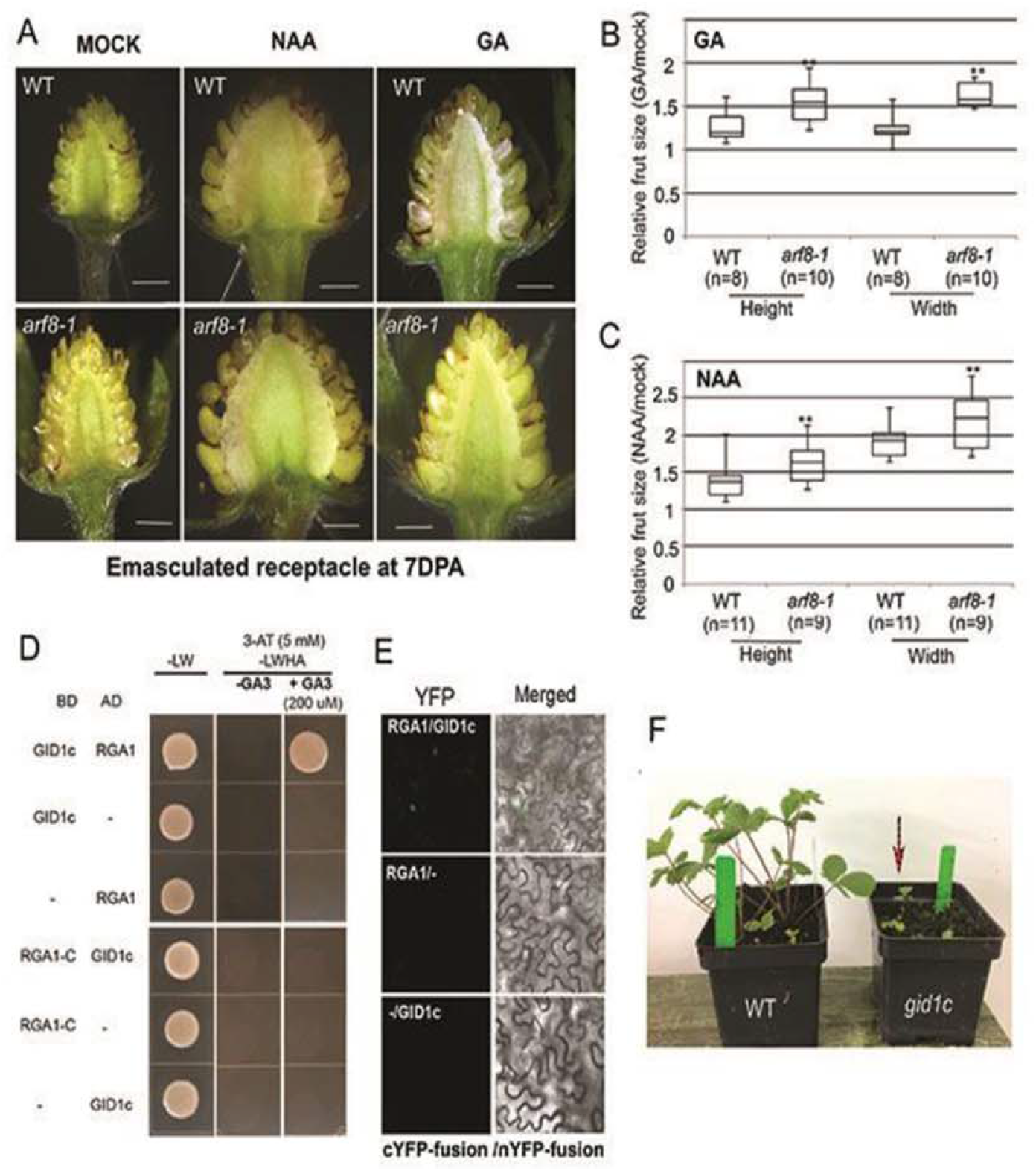
*arf8-1* exhibits enhanced response to GA and auxin. A. Photos of bisected stage 3 receptacles of WT and *arf8-1.* The flowers were first emasculated and then applied with NAA, GA3, or mock solution (see Methods) until 7DPA. Bars; 1 mm. B. Relative receptacle size of WT and *arf8-1* at stage 3 (7DPA) after GA treatment of emasculated flowers. The height and width of hormone GA-treated receptacles are divided respectively by the height and width of mock treated receptacles of the same genotype. ** above bar represents significant difference (P < 0.01) between WT and *arf8-1*. C. Relative receptacle size of WT and *arf8-1* at stage 3 (7DPA) after auxin (NAA) treatment of emasculated flowers. Data is similarly measured and analyzed as B. D. Yeast two-hybrid assay showing the interaction between FveGID1c and FveRGA1 only in the presence of GA3 (200 μM). The interaction is dependent on the DELLA domain, which, when deleted in RGA1-C, could no longer support the interaction even in the presence of GA. E. BIFC assay in tobacco leaf cells confirming positive interaction between FveRGA1 and FveGID1c. The tobacco cells may produce GA to enable the interaction. F. A CRISPR/CAS9-induced *gid1c (−149, −166+15)* biallelic mutant exhibiting severe dwarf.

While it is understandable that *arf8-1,* a mutant gene in the auxin signaling pathway, affects sensitivity or response to auxin, why does *arf8-1* also exhibit increased sensitivity or response to GA? One possibility is that *FveARF8* regulates the expression of GA receptors to affect the sensitivity to GA. Three *FveGID1* genes were identified from the *F. vesca* genome (gene22353, gene20092, and gene27756) as the candidates of GA receptor (Figure S6), only *gene22353,* named as either *FveGID1a* (Kang et al., 2013) or *FveGID1c* (Li et al., 2019) and thereafter, is specifically expressed in the receptacle at early fruit stages (Figure S6B-D). If *FveGID1c* encodes a GA receptor for fruit development, FveGID1c should bind FveRGA1 when GA is present. Indeed, FveGID1c directly interacts with FveRGA1 in yeast only when GA is added to the medium (Figure 4D) and this interaction requires the DELLA domain at the N-terminus as FveRGA1-C (missing N-terminal 564 bp) fails to interact with FveGID1c even when GA is added (Figure 4D). The interaction between FvGID1c and FveRGA1 is also demonstrated in tobacco cells via BiFC (Figure 4E). CRISPR/Cas9 was used to knockout GID1c; a biallelic mutant *gid1c (−149; −166+15)* was obtained (Figure S1C), which exhibited strong dwarf but never flowered (Figure 4F). Although the GID1c sgRNA1 and sgRNA2 show 5 and 9 mismatches respectively to the other two GID1 genes (gene20092 and gene27756) (Figure S6E), we still sequenced the other two GID1 genes (gene20092 and gene27756) from the *gid1c (−149; −166+15)* plant and found no mutations in them. The strong dwarf phenotype indicates that *FveGID1c* encodes an important GA receptor.

Next, we tested if *FveGID1c* expression is altered in *arf8-1* at the early fruit stages 1-3. *FveGID1c* expression level is similar in *arf8-1* and WT at stage 1; at stages 2-3, *FveGID1c* expression is reduced in WT but remains high in *arf8-1* mutant receptacle, significantly higher in *arf8-1* when compared with WT (Figure 5), indicating that *FveARF8* may limit the sensitivity to GA by down-regulating *FveGID1c* at stages 2-3. Given that FveARF8 is no longer inhibited by FveRGA1 at stages 2-3 due to fertilization-induced GA production and subsequent FveRGA degradation, FveARF8 may then be free to repress *FveGID1c* expression at stage 2-3 to dampen sensitivity to GA in feedback regulation.

**Figure 5.**
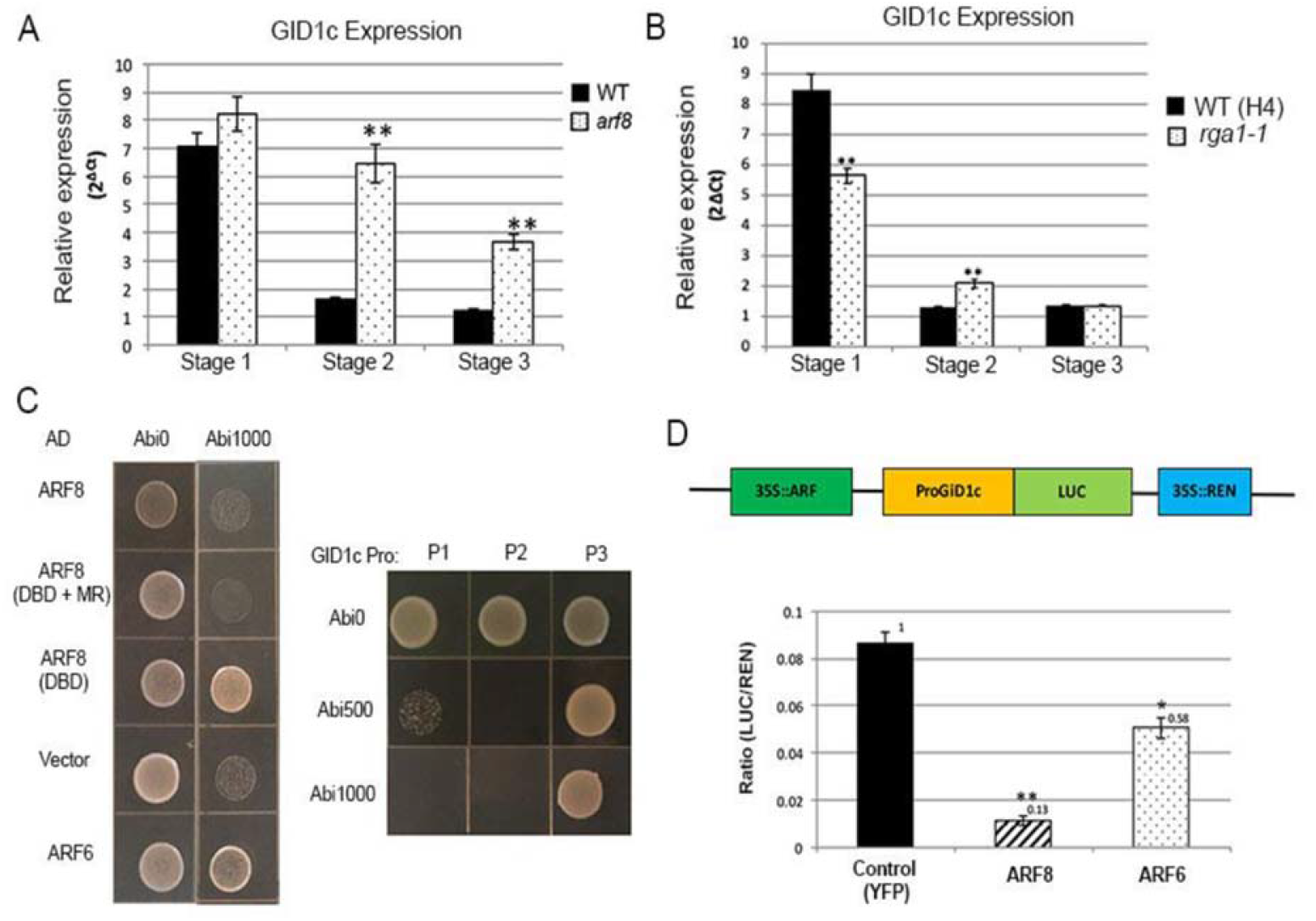
FveARF8 and FveARF6 directly represses the expression of *FveGID1c*. A. qRT-PCR showing relative expression of *FveGID1c* in WT (YW5AF7) and *arf8-1* mutants at stage 1-3. B. qRT-PCR showing relative expression of *FveGID1c* in WT (H4) and *rga1-1* mutants at stage 1-3. C. Y1H assay showing activation of GID1c Promote fragment 2 (P2) when FveARF6 or ARF8-DBD domain fused to GAL4 AD is introduced into the yeast. On the right, three different promoter fragments (P1, P2, and P3) are tested for self-activation at two different concentrations of antibiotics Abi500 and Abi1000. P2 does not have any self-activation activity. D. Diagram of the LUC reporter system. The expression of LUC driven by the GID1c promoter is repressed strongly by *35S::FveARF8* and weakly by *35S::FveARF6.* Robust self-activation in the absence of transacting factor (with YFP serving as a negative control) is shown when 1 μm GA_3_ is added in all essays. Y-axis shows relative LUC expression normalized against 35S::REN expressed from the same vector.

We then tested *FveGID1c* transcript level in *rga1-1* mutant background. In *rga1-1* mutant, a reduction of *FveGID1c* transcript is observed at stage 1 (Figure 5B), consistent with that de-repressed *FveARF8* in *rga1-1* could act to repress *FveGID1c* transcription. Given that *FveARF8’s* function may still be constrained by IAAs at stage 1, the repression of *FveGID1c* is moderate (Figure 5B). At stage 2-3 (post-fertilization), freed from inhibition by FveIAAs and FveRGA1, FveARF8 can repress *FveGID1c* expression in WT as well as in *rga1-1* (Figure 5B). The above qRT-PCR data are consistent with a negative regulatory relationship of *FveRGA1* to *FveARF8* and to *FveGID1c.*

To test if FveARF8 directly represses *FveGID1c,* three segments of *FveGID1c* promoters, P1, P2, and P3 (Figure S6F), were respectively fused to the AUR1-C reporter and integrated into the yeast strain in a Y1H system. Self-activating activity of each segment was examined at two Abi concentrations, 500 μM and 1000 μM (Figure 5C). Promoter fragment 2 (P2), which does not self-activate and contains two potential AuxREs and two potential G-boxes (Figure S6F), was used in the Y1H against transcription factor FveARF6-AD or FveARF8-AD (Figure 5C). FveARF6-AD and FveARF8 (DBD)-AD readily activate the *FveGID1c* (P2)::AUR1-C reporter, indicating direct binding of FveARF8 and FveARF6 to the *FveGID1c* promoter. However, full length FveARF8 or a truncated FveARF8 (DBD + MR) fail to activate the *FveGID1c* (P2) reporter (Figure 5C), indicating that the MR domain of FveARF8 may possess strong repressor activity that could counter Gal4-AD.

A transient expression reporter system in tobacco cells is also used to test direct effect of FveARF8 and FveARF6 on the *FveGID1c* promoter. A 1kb promoter fragment of *FveGID1c* was fused to the firefly luciferase (LUC) and integrated into vector PLAH-LARM (Taylor-Teeples et al., 2015), which also contains *35S::REN (Renilla luciferase)* as a control and *35S::FveARF8* or *35S::FveARF6* (Figure 5D). 1 μM GA was applied to the tobacco cells to cause robust expression from *pGID1c::LUC* perhaps due to endogenous transcription factors mediating GA responses. Compared with *35S::YFP* control; *35S::FveARF8* significantly reduces the expression from *pGID1c::LUC* (Figure 5D), suggesting that FveARF8 harbors strong repressor activity and binds directly to the promoter of *FveGID1c.* FveARF6 appears to possess very weak repressor activity (Figure 5D). Together, these data suggest that FveARF8, and perhaps FveARF6, may directly bind and repress *FveGID1c* to limit sensitivity to GA at post-fertilization stage.

### Consensus co-expression network identifies *FveARF8, FveARF6,* and *FveIAA4* in the same co-expression module

Our data revealed a previously unknown mechanism of GA-auxin crosstalk where auxin represses the sensitivity to GA via activation of *FveARF8.* In auxin signaling pathway, AUX/IAA proteins are known to bind and repress ARFs in the absence of auxin (Guilfoyle, 2015; Salehin et al., 2015). Which *FveIAAs* could bind and inhibit *FveARF8? F. vesca* genome V4.0a2 has about 34 *FveARFs* and 25 *FveIAAs* (Edger et al., 2018; Li et al., 2019). To identify candidate *FveIAAs,* we mined the consensus co-expression network generated previously (Shahan et al., 2018) (http://159.203.72.198:3838/fvesca/). *FveARF8* resides in module 80 of the Consensus_90 Fruit Network; this module 80 correlates positively in expression with early stage receptacle fruit tissues. Interestingly, the two enriched GO terms for this module are “auxin activating signaling pathway” and “cellular response to auxin stimulus”. *FveARF6* and *FveIAA4* are found in this module with a high correlation co-efficient (>0.91) with *FveARF8* (Figure 6; Data S1). Based on RNA-seq data (Kang et al., 2013; Hollender et al., 2014; Hawkins et al., 2017), all three genes are highly and specifically expressed in the receptacle, ovary wall and style during early fruit development at stage 1-5 (Figure 6B). To validate their expression, qRT-PCR was conducted using receptacle at fruit stages 1, 2, and 3; all three genes are expressed in the receptacle at stage 1 (pre-fertilization) and show downward expression trend post-fertilization at stage 2 and 3 (Figure 6C).

**Figure 6.**
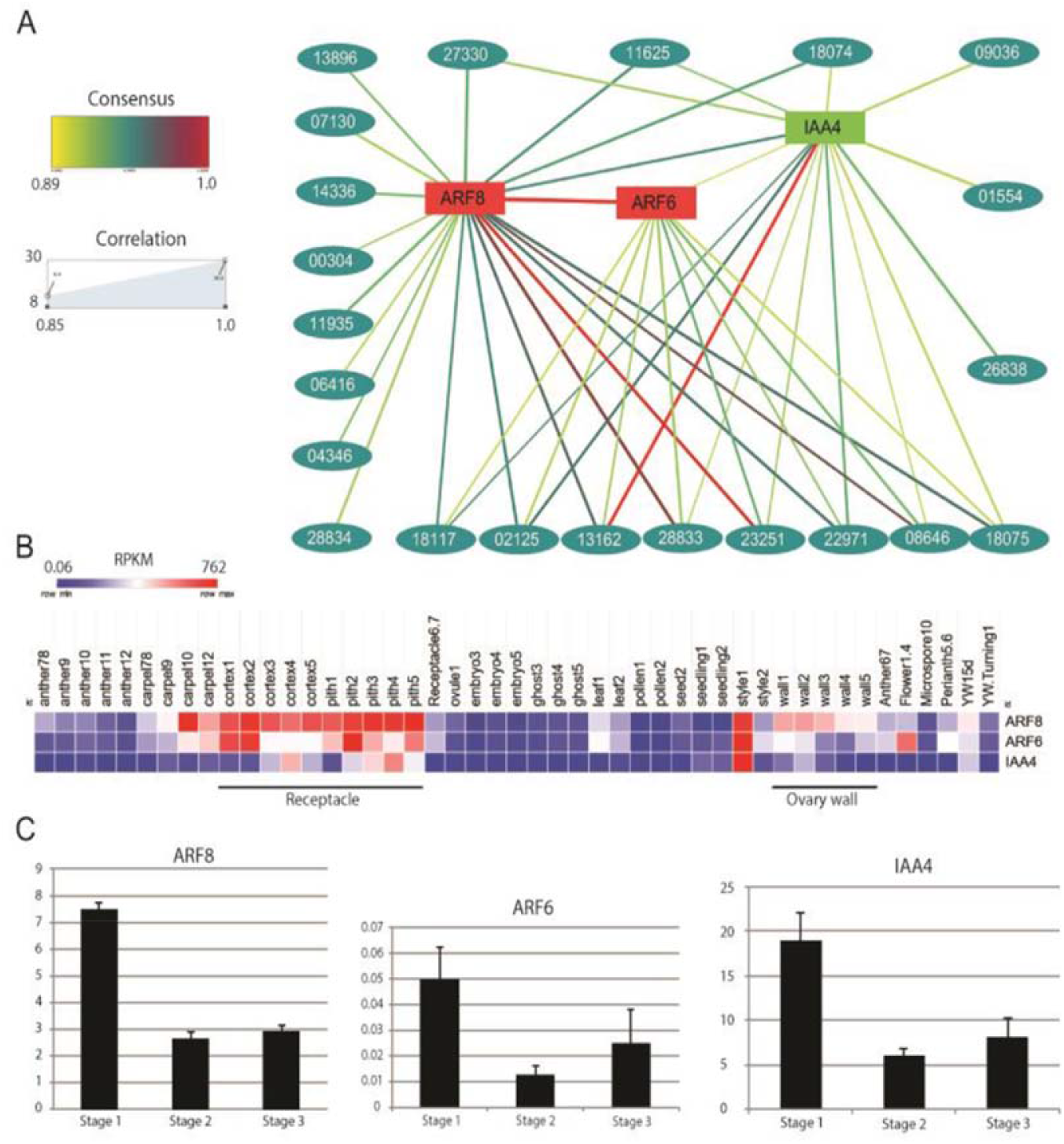
Network analysis identifies *FveARF8, FveARF6* and *FveIAA4* as potential regulators of fruit initiation. A. Consensus Network analysis identifies *FveARF8, FveARF6,* and *FveIAA4* in module 80 of Consensus 90 Fruit Network. The five-digit ID numbers for all other genes in the same module are shown and also listed in Table S1. This module correlates with young receptacle fruit. The consensus score and correlation score between genes are indicated by color and thickness of lines respectively. B. Heatmap of *FveARF6, FveARF8* and *FveIAA4* based on prior RNA-seq data, revealing specific expression in young fruit and style. C. qRT-PCR of *FveARF8, FveARF6,* and *FveIAA4* in developing receptacle (pith and cortex) at stages 1-3. Data represents mean ± SE of gene expression from three biological replicates of YW receptacle tissues. The Y-axis indicates relative expression level compared to reference gene PP2a (gene03773) derived using formula 2^-ΔCt^. Different assaying techniques and tissue sampling methods may contribute to some differences between qRT-PCR result and the RNA-seq data in (B).

To test if FveIAA4 in the same network module may interact and repress *FveARF6/8* in the absence of auxin, Y2H shows that FveIAA4 interacts with both FveARF8 and FveARF6 (Figure 7). The C-terminal PB1 domain of FveARF8 is shown to be both necessary and sufficient for the interaction with FveIAA4 (Figure 7B). BiFC confirms these interactions (Figure 7C). The data suggest that fertilization-induced auxin may cause degradation of FveIAA4, releasing FveARF8/6 to exert positive or negative regulatory effects on downstream genes.

**Figure 7.**
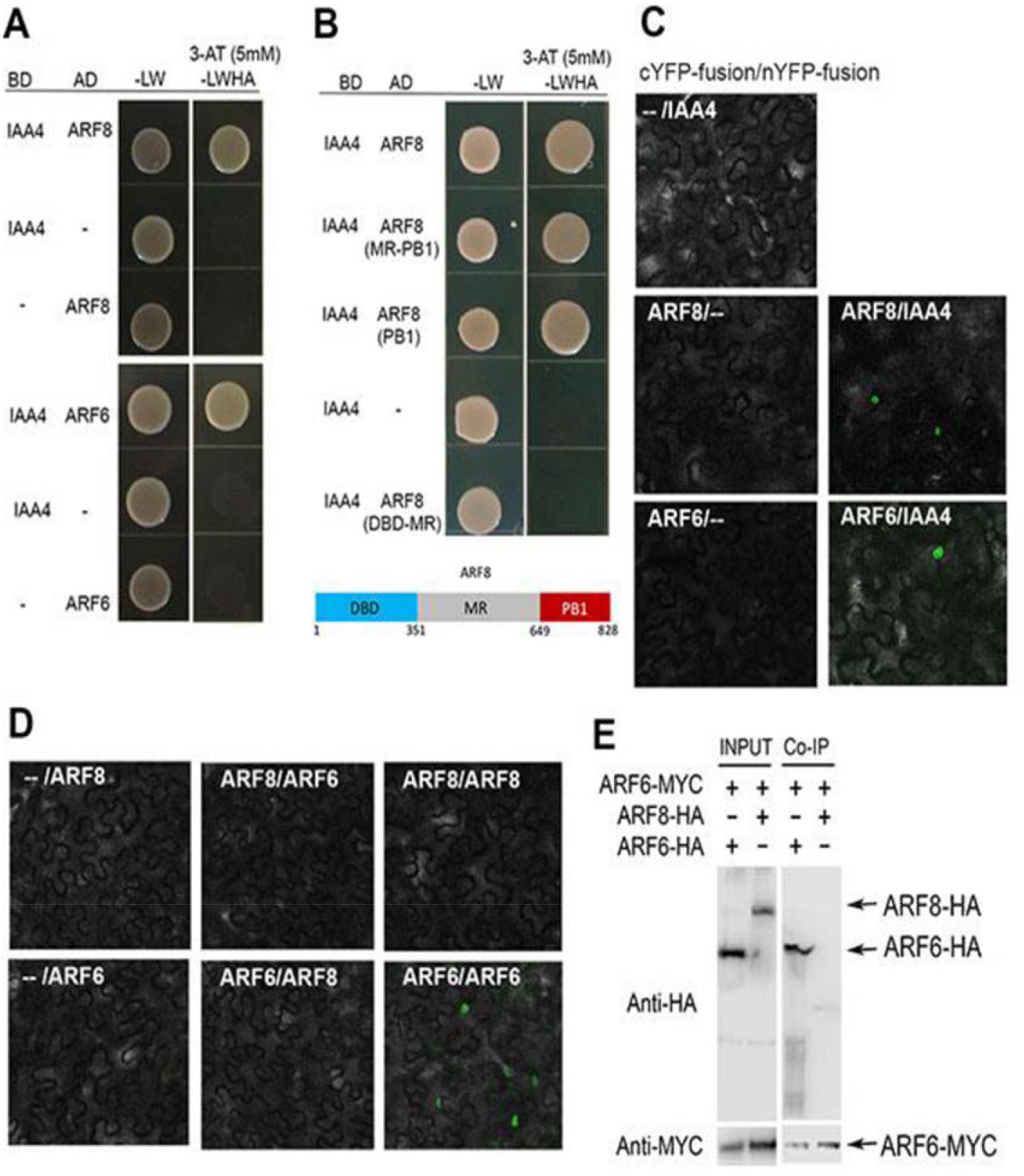
Yeast two-hybrid, BiFC, and co-immunoprecipitation assays testing the interaction between FveIAA4 and FveARF8/FveARF6. A. Yeast two-hybrid assay revealing the positive interaction between FveIAA4 and FveARF6/FveARF8. B. Domain dissection of FveARF8 showing that the PB1 domain is necessary and sufficient for the interaction with FveIAA4. A diagram of ARF8 is drawn beneath, showing DNA binding (DBD), Middle Region (MR), and PB1 domains. Numbers beneath the diagram indicate amino acid residues. C. FveARF8 and FveARF6 respectively interact with FveIAA4 shown by BiFC in tobacco cells. D. BiFC in tobacco cells showing a lack of interaction between FveARF8 and FveARF6. While FveARF8 does not homodimerize, FveARF6 does. E. Co-immunoprecipitation using protein extracts from yeast cells containing c-Myc-tagged Ga14BD-FveARF6 and HA-tagged Ga14AD fusion constructs. INPUT lanes show western blots of yeast protein extracts containing indicated constructs. Co-IP lanes are western blots of immunoprecipitated proteins with anti-HA antibody.

To test if FveARF6 and FveARF8 could work together as heterodimers, we first performed BiFC (Figure 7D) which showed that while FveARF6 and FveARF8 do not interact with each other, FveARF6, but not FveARF8, interacts with itself (Figure 7D). FveARF8’s inability to interact with FveARF6 is subsequently confirmed in coimmunoprecipitation assay (Figure 7E) using protein extracts from yeast containing MYC-tagged Ga14BD-FveARF6 and HA-tagged Ga14AD-FveARF8 or Ga14AD-FveARF6.

The different MR sequence between FveARF8 and FveARF6 protein (Figure S5) combined with their distinct ability to hormodimerize suggests that *FveARF6* and *FveARF8* may not be functionally redundant. CRISPR construct was made for *FveARF6* and transformed into WT *F. vesca* as well as *arf8-1 (−2, −2) line# 156 that has segregated away the original CRISPR-sgARF8 transgene*. While we failed to obtain an *arf6* single mutant, *arf6-1 (−1, −1)* mutation was identified in *arf8-1 (−2, −2)* background in two different transgenic lines (Figure S1D). These *arf6-1; arf8-1* double mutant lines developed fruits similar in size to WT and significantly smaller than *arf8-1* single mutants (Figure S7). The phenotype supports that *FveARF6* and *FveARF8* may play different roles during fruit initiation and enlargement.

## Discussion

Strawberry has long been a model to study fruit development, especially fruit set, when Nitsch (1950) first observed the effect of auxin in stimulating receptacle fruit fleshy formation when achenes were removed (Nitsch, 1950). However, the molecular mechanisms underlying this elegant communication between seed and fruit flesh are not well understood. Further, being the accessory fruit, where the fruit fleshy is derived from receptacle instead of ovary wall, it poses additional questions concerning if the conserved fertilization signals, auxin and GA, activate downstream fruit program in a similar or distinct mechanism from botanical fruits. The work presented here provides basic molecular frame work on how the GA and auxin signaling components interact and coordinate fleshy fruit program in strawberry receptacle using both conserved and novel mechanisms. The work laid the foundation for future engineering of parthenocarpic strawberry for enhancing fruit productivity when pollinators are scarce.

### *FveRGA1* is a key regulator of fruit set in diploid strawberry, but *FveARF8* is involved in downstream regulation of fruit development post-fertilization

*rga1-1* is the only strawberry mutant so far reported to develop parthenocarpic fruit, indicating *FveRGA1* as a key regulator of fruit set in strawberry. FveRGA1 normally prevents fruit sets by inhibition of a large number of TFs through protein-protein interaction. In *rga1-1* mutants, the inhibition is absent, and the TFs are free to activate downstream genes for fruit development even in the absence of plant hormone, leading to parthenocarpic fruit. This mutational effect of *rga1-1* mimics the application of GA in emasculated flowers, where GA-GID1c induced degradation of FveRGA1 lead to parthenocarpic fruit (Kang et al., 2013). Our study identifies *FveRGA1* as a target gene for genetic manipulation to produce parthenocarpic strawberry.

The larger fruit size of *arf8* mutants appears caused by different reasons from *rga1-1.* Further, the effect of *arf8* on receptacle size can be divided into pre- and post-fertilization stages. At the pre-fertilization stage, the larger receptacle size is likely due to a defect on flower development, which is also reflected by increased petal numbers in *arf8* mutants (Figure S2). It is important to note that the *arf8-1* mutant receptacle size stays the same from stage 1 to stage 3 when fertilization is prevented (Figure 2A-B). In contrast, at the post-fertilization stages when auxin and GA are produced, the faster increase in fruit size of *arf8-1* mutants reflects an enhanced sensitivity to auxin and GA (Figure 4A-C). In other words, same concentrations of phytohormones could mount a stronger response in *arf8-1* mutants. This is why *arf8-1* mutants do not develop parthenocarpic fruit as GA and auxin are still required for the *arf8-1* mutants to develop fruit.

### A model on the mechanism of GA-auxin crosstalk on fruit initiation

Figure 8 is a model that both summarizes previous understanding of GA and auxin signaling pathways and incorporates our findings here. First, auxin and GA signaling pathways act in parallel to promote fruit set and fruit development (thick green curved arrows). At pre-fertilization (stage 1) receptacle (Figure 8B), RGA1 and IAA4, respectively bind and inhibits a large number of TFs including ARF8 (red bars), blocking fruit development. At post-fertilization (stage 2) receptacle (Figure 8C), GA and auxin are produced from the fertilized seed and transported to the receptacle, where GA and auxin interact with their respective receptors, FveGID1c and FveTIR, which subsequently facilitate ubiquitin-mediated degradation of FveRGA1 and FveIAA4. Therefore, upon fertilization, TFs in auxin and GA pathways are activated to promote fruit set and subsequent fruit development (blue arrows).

**Figure 8.**
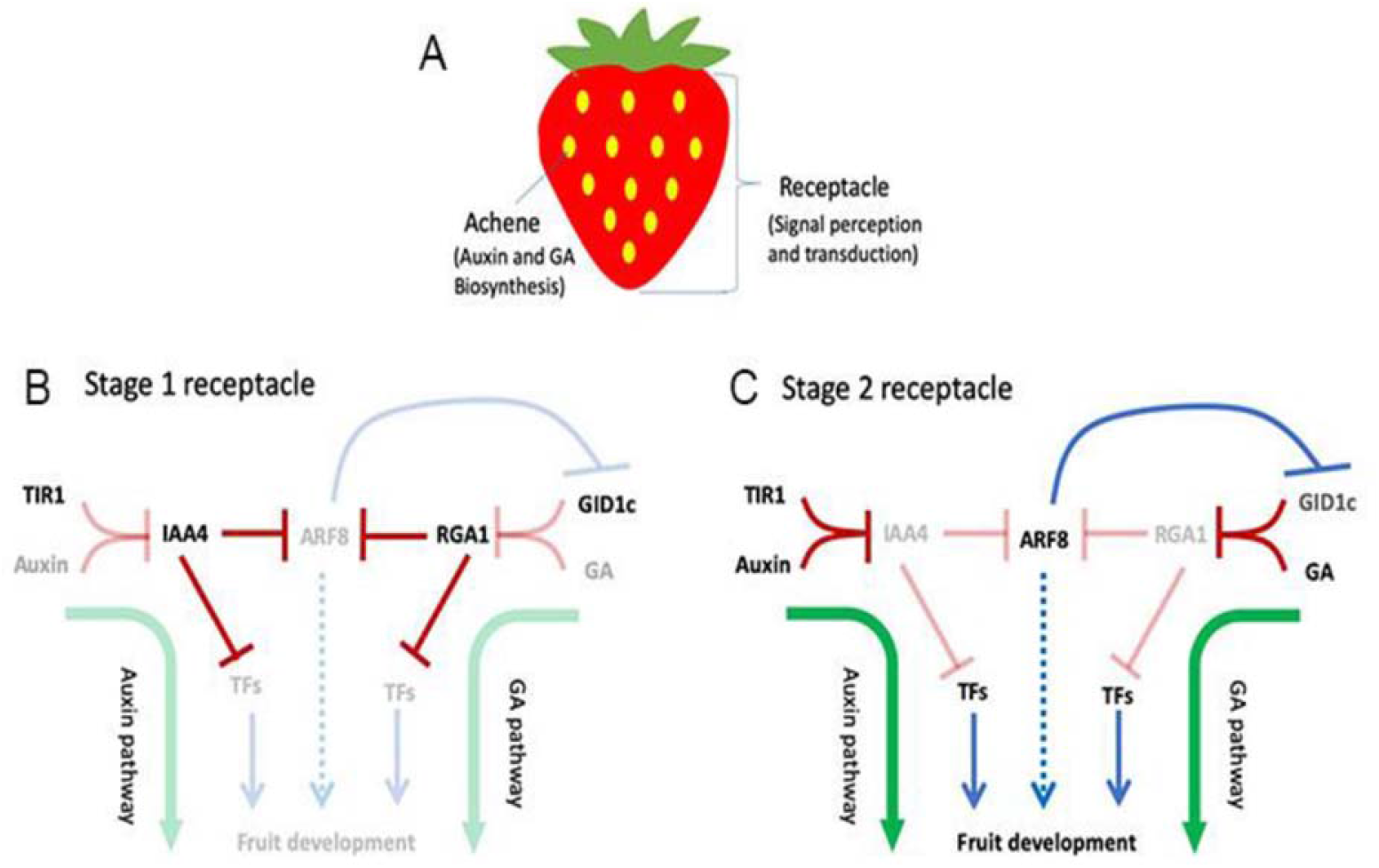
A model that summarizes auxin/GA crosstalk in strawberry fruit initiation. A. A diagram illustrating spatial separation of hormone synthesis in the achene from hormonal signal perception and transduction in the receptacle. B. In receptacle at stage 1 (pre-fertilization), when auxin and GA are absent, IAA4 and RGA1 bind and repress ARF8 and other TFs in the auxin and GA signaling pathway. Fruit set is blocked. C. In the receptacle at stage 2 (post-fertilization), auxin and GA are transported to the receptacle (from achenes) to induce signal transduction. Specifically, GA/GID1c and auxin/TIR receptor complex caused degradation of FveRGA1 and FveIAA4, releasing FveARF8 and TFs which promote fruit set and subsequent development. FveARF8 at the same time represses the transcription of FveGID1c in a feedback regulation. In B and C, the red lines describe negative regulation at post-translational level via protein degradation or binding of repressors to transcription factors (TFs). Blue lines describe positive (arrows) or negative (bar) regulation at transcription level. Light shaded lines or names indicate “inactive” or “absent”. Dotted blue line from *FveARF8* indicates *FveARF8’s* putative and indirect role in promoting fruit development.

We show that FveARF8’s DBD-MR and PB1 domains interact respectively with FveRGA1 and FveIAA4 (Figure 3 and 7), indicating that FveARF8 activity could be simultaneously inhibited by FveIAA4 and FveRGA1 prior to fertilization (Figure 8B). At stage 2, when both auxin and GA are produced and FveIAA4 and FveRGA1 are both degraded, FveARF8 and other TFs are now freed from both inhibitions to regulate the transcription of downstream genes (blue lines) (Figure 8C). This explains why combined treatment of auxin and GA is required to cause emasculated fruit to develop to the same size as pollinated fruit (Kang et al., 2013).

One of the downstream targets of *FveARF8* is the *FveGID1c* gene, and FveARF8 represses *FveGID1c* transcription to dampen sensitivity to GA in a feedback regulatory loop (Figure 8C). This transcriptional down-regulation of *FveGID1c* by *FveARF8* at stage 2 is detected by qRT-PCR (Figure 5A) and appears to be direct (Figure 5C-D). As ARFs normally have a large number of target genes in a genome (Oh et al., 2014), *FveARF8* may also promote fruit development indirectly through repressing other negative regulators of fruit enlargement. This is indicated by a dotted blue arrow from *FveARF8* in Figure 8C.

### *FveGID1c (gene22353)* encodes a GA receptor in *F. vesca*

Previous gene expression analysis (Kang et al., 2013), current Y2H assay in the presence or absence of GA (Figure 4D), and CRISPR/CAS9-induced knockout of *FveGID1c* (Figure 4F) all support that *FveGID1c* is an important GA receptor. While there are three *FveGID1* genes (Figure S6A), *FveGID1c (gene22353)* not only exhibits significantly higher expression than the other two *FveGID1b* genes (gene20092 and gene27756) but also is more specifically expressed in the receptacle (Figure S6B-D). The severe phenotype of *gid1c (−149, −166+15)* prevented the mutant plant to produce flowers. Future work using inducible RNAi maybe necessary to allow the examination of *gid1c* phenotype in fruit development.

### Both conserved and novel GA-auxin crosstalk mechanisms exist in strawberry

While both auxin and GA act to promote fruit set in several organisms including Arabidopsis, tomato, and strawberry, their interaction and crosstalk mechanism may vary. In the botanical fruit of *Arabidopsis* and tomato, auxin was shown to act upstream of GA by promoting GA biosynthesis (Serrani et al., 2008; Dorcey et al., 2009; Fuentes et al., 2012). Recent work in tomato revealed another crosstalk point through direct proteinprotein interactions between SlARF7 and SlDELLA as well as SlIAA9 via SlARF7’s MR2 and PB1 domains respectively (Hu et al., 2018). However, it is not known if strawberry employs similar or novel crosstalk mechanisms to induce fruit formation given the distinct floral tissue used by strawberry to form fruit flesh.

Our finding of direct interaction of FveIAA4 with FveARF8/FveARF6 and of RGA1 with FveARF8/FveARF6 illustrates conservation of crosstalk mechanism in a different spatial context (ie. in accessary fruit). However, our data also reveal a previously unknown crosstalk mechanism where FveARF8 represses *FveGID1c* transcription to reduce sensitivity to GA in a feedback regulatory loop (Figure 8). The enhanced sensitivity to auxin in *arf8-1* (Figure 4A) also suggests that *FveARF8* may act in a feedback loop of auxin signaling. Proper regulation of FveARF8 protein activity through FveIAA4 and FveRGA1 may serve to ensure that auxin and GA are limiting each other’s signaling pathway to balance each other’s impact on fruit enlargement. One interesting outcome of this balance is reflected in fruit shape. Previously, exogenous application of GA to *F. vesca* fruit was shown to impact fruit shape differently from exogenous application of NAA (Liao et al., 2018). Similarly, we showed that treatment of emasculated WT or *arf8-1* with NAA significantly increases fruit width and hence rounder fruit shape, while treatment with GA significantly increases fruit length and hence longer fruit shape (Figure 4A). Hence, a balance between the two hormonal pathways may impact fruit shape as well as size.

### FveARF8 and FveARF6 exhibit differences in protein sequence, dimerization, repression strength, and phenotype

Most ARF proteins can be divided into three domains, a N-terminal domain for DNA-binding and dimerization, a variable middle region (MR) that confers activation or repression activity, and a C-terminal PB1 domain for interaction with ARFs as well as Aux/IAA inhibitors (Tiwari et al., 2003; Boer et al., 2014; Guilfoyle, 2015; Powers et al., 2019). A recent report indicates that Arabidopsis activator ARF7 and ARF19 form cytoplasmic assemblies as a mechanism to sequester them to modulate cellular competence to respond to auxin (Powers et al., 2019). Sequence divergence in all three domains contribute to distinct functional properties of ARFs within and between species.

Arabidopsis *ARF6* and *ARF8* genes are classified as activator ARFs that promote both vegetative and reproductive development (Ulmasov et al., 1999; Tiwari et al., 2003). Arabidopsis *arf6; arf8* double mutants show more severe floral phenotypes than either single mutants, suggesting partial redundancy between *AtARF6* and AtARF8 (Nagpal et al., 2005). In Arabidopsis and tomato, over-expression of *miR167,* which targets both ARF6 and ARF8, also caused similar phenotypes as the *arf6; arf8* double mutants (Wu et al., 2006; Liu et al., 2014; Zheng et al., 2019). Most relevant to this work, *Arabidopsis arf8* mutants develop parthenocarpic fruit, and the expression of an “dominant negative” *AtARF8* causes parthenocarpy in Arabidopsis and tomato (Goetz et al., 2007). Thus, ARF8 possesses a conserved role in preventing fruit set in Arabidopsis and tomato.

In this study, *FveARF8 (gene31631)* and *FveARF6 (gene30394)* are identified and named based on their blast best hit with their Arabidopsis counterparts. A second *F. vesca ARF6* genes (*gene22728*) is not studied here since it is not in the same co-expression module as *FveARF8.* Sequence alignment among AtARF6/AtARF8/FveARF6/FveARF8 reveals high levels of sequence conservation at the N-terminal DBD and C-terminal PB1 domains but poor sequence similarity at the MR domain (Figure S5). When compared with the MR of AtARF6, MR of FveARF6 has a large 40 amino acid deletion and a reduced number of polyQ tracks (Figure S5), which may weaken the activation function of FveARF6. Together, the poor sequence conservation in the MR domain, different strength of repression on *FveGID1c* promoter, and different ability to homodimerize suggest that *FveARF6* and *FveARF8* may function differently. This is supported by the fruit phenotypes where a loss-of-*FveARF6* led to fruit size reduction in *arf8-1* background and a loss-of-*FveARF8* caused fruit size increase (Figure S7).

### Consensus co-expression network is a powerful tool

Previously, a consensus-clustering approach was applied to the standard WGCNA (Weighted Gene Co-expression Network Analysis) to generate robust and reproducible modules (Shahan et al., 2018). Since ARF and IAA genes are from large gene families, the consensus co-expression network led us to a co-expression module consisting of *FveIAA4, FveARF6* and *FveARF8* (Figure 6A). Our results support their protein-protein interaction and functional roles in early fruit development and demonstrate the predictive power of consensus co-expression network analysis. Such network is especially valuable in guiding selection of genes belonging to large gene families as well as studies in non-model species where transformation is resource-intensive and challenging.

## Methods

### Plant materials and growth condition

The *Fragaria vesca* cultivar Yellow Wonder 5AF7 (YW5AF7) was used in most of this study. Cultivar Hawaii 4 (H4) was used in generating BZR1-RNAi lines. Plants were grown in a growth chamber at a temperature of 25 °C/16 hour light followed by 22 °C/8 hour darkness with a relative humidity of 50%. The *F. vesca* genes studied here are indicated by their gene IDs (genome version 2.0.a2; genome version 4.0.a2): *FveARF8 (gene31631; FvH4_1g27650), FveARF6 (gene30394; FvH4_3g05010), FveIAA4 (gene 16569; FvH4_6g02870), FveGID1c (gene22353; FvH4_6g04960),* and *FveRGA1 (gene 06210; FvH4_4g34110).* The names *FveARF8, FveARF6, and FveIAA4* are assigned based on their best blast hit with their Arabidopsis counterparts.

*arf8* mutants were generated using CRISPR/CAS9 (Zhou et al., 2018). Four independent *arf8* transgenic lines #100 (−2, −2), #156 (−2, −2), #391-3 (−8, −23+5), and #391-31 (−2+1, − 2+1) were detailed in Figure S1. Line #100 (−2, −2) and line #156 (−2, −2) are referred to as *arf8-1* in the text. T1 generation *arf8* mutant plants were analyzed, and about 20% of them have segregated away the CRISPR-sgARF8 transgene.

The *rga1-1* (u230) mutant was previously generated by ENU chemical mutagenesis in a *ga20ox4* loss of function mutant background (Tenreira et al., 2017; Caruana et al., 2018). This *rga1-1; ga20ox4* was backcrossed with WT (H4) to yield *rga1-1* single mutant, which was used in all analyses. H4 was used as WT control whenever *rga1-1* was used.

### CRISPR/Cas9 genome editing and over-expression

To generate the *arf8; arf6* double mutants, two seed gRNA sites (gRNA1: TGCTTCAACCAACATGGAAG and gRNA2: GCCAGTGACACTAGTACCCA) targeting an *FveARF6 (gene30394)* exon were inserted into the JH4 entry vector and then incorporated into binary vector JH19 via gateway cloning (Zhou et al., 2018). Cotyledon of *arf8* mutant line #156 (−2, −2), which has segregated away its prior sgARF8-Cas9 construct, was used for transformation with the CRISPR-sgARF6 construct. Four transgenic lines were generated. To detect mutation, genomic sequences spanning two target sites of *FveARF6* were amplified by primers FvARF6-F5 and FvARF6-R5 (Data S2) and analyzed by Sanger sequencing.

The *FveGID1c* CRISPR construct was created using JH4 and JH17 vectors (Zhou et al., 2018). Two seed guide RNA sequences (gRNA1: GCAACTGAGACACGCCTGG and gRNA2: GTTCCCAAGGAGCCTTGTGG) targeting *FveGID1c* were inserted into the entry vector JH4 and incorporated into binary vector JH17 via gateway cloning. In total, 3 transgenic lines were generated. The genomic region spanning the two target sites in *FveGID1c* was amplified by primers FvGID1a-F1 and FvGID1a-R1 (Data S2) and sequenced. Only one transgenic line, line #7 *gid1c (−149, −166+15),* has a clear bi-allelic mutation (Figure 4F).

To test if the other two *FveGiD1b* genes (gene20092 and gene27756) were also mutated in *gid1c* line #7, primer pairs gene20092-F&gene20092-R and gene27756-F&gene27756-R (Data S2) were used to respectively amplify the genome sequence of the two *FveGiD1b* genes. The PCR products were then sequenced.

To generate the FveARF8 over-expression (OX) construct, full-length *FveARF8* CDS was PCR amplified from YW5AF7cDNA library with primers FvARF8-SalI-F and FvARF8-Kpn-R (Data S2) and inserted into pENTR-2b vector via SalI/KpnI. The pENTR-FveARF8 was incorporated into JH23 destination vector through gateway cloning. In total, eight over-expression transgenic lines were generated and analyzed.

To construct the over-expression vector JH23 (Figure S6), the OCS terminator was amplified from JH19 (Zhou et al., 2018) and inserted into pMDC99(Curtis and Grossniklaus, 2003) at PacI and SacI sites; the AtUBQ10 promoter was amplified from JH19 (Zhou et al., 2018) and then inserted into above PMDC99-OCS at HindIII and KpnI sites to yield JH23.

Most above constructs were transformed in *F. vesca* (YW5AF7 or *arf8-1* mutant in YW5AF7 background) using previously published protocols (Kang et al., 2013; Caruana et al., 2018).

### Receptacle size measurement

The receptacle of *arf8* mutant and WT (YW5AF7 or Hawaii 4) were cut in half and imaged under Stemi SV 6 dissecting microscope (Zeiss. Inc). The height and width were then measured using the ZEN software. Flowers that just open with anthers not yet dehiscent were considered stage 1 (0DPA). Flowers at 5-7 DPA were considered stage 3 usually with only 1 petal left. Receptacle measurements were normalized to petal size (height and width of receptacle divided respectively by height and width of petal) to yield relative receptacle size in Figure 1. To measure the receptacle size of *rga1-1* (Figure 2) the newly opened but non-dehiscent flowers were labeled as 1DPA (stage 1), the receptacle length and width were measured at 9 DPA (stage 4) and 11-12 DPA (stage 5). To determine parthenocarpy, *arf8-1* or *rga1-1* flowers were emasculated to prevent self-fertilization. Fruit size at 0DPA (pre-fertilization) was compared to fruit size at 7 DPA (post-emasculation) within the same genotype.

For measuring the receptacle size of GA3- and NAA-treated fruits, WT (YW5AF7) or *arf8* flowers were emasculated at 0DPA under Stemi SV 6 dissecting microscope (Zeiss. Inc), 1mM GA3 or 1mM NAA mixed with 0.1% Tween 20 was applied to the emasculated receptacle. This treatment was repeated every two days until day 7 when the receptacle size was measured. The relative GA- and NAA-treated receptacle height and width were normalized to mock treated fruit of the same genotype.

For *arf8-1,* 8-19 fruits derived from 5 to 10 plants of the same genotype were measured; and then combined to yield one bar in bar graph (or one box in box plot). For YW5AF7 (WT) control, fruit measurement was similarly made on fruits from 12 plants. Measurement of other *arf8* CRISPR lines were similarly conducted. For hormone treatment experiments, the same groups of *arf8-1* and YW5AF7 plants above were used in fruit measurement. For *rga1-1*, twelve mutant plants were grown from the same original mother plant through vegetative propagation via runner. Ten H4 (WT) control plants were similarly propagated through an original H4 mother plant. The same measurement and analysis strategy described for *arf8-1* was used. For *arf8-1*; *arf6-1* double mutants, two transgenic lines were used, and each line has 3-4 plants derived from the same calli. 10 YW5AF7 (WT) plants were grown together and used as controls.

### Statistical analysis

Data from fruit size measurements, qRT-PCR, and LUC assay were all analyzed using unpaired, unequal variance, two-tailed student’s t test. Significant difference (P <0.01) and difference (P <0.05) between treatments or genotypes are denoted by **and *, respectively. Data are expressed as the means ± standard deviation for different biological replicates.

### BiFC and Y2H to test protein-protein interaction

For the BiFC, full-length CDS of *FveARF8, FveARF6* were PCR amplified from cDNA and ligated into PXY105 (cYFP) at BamHI and SalI sites, full-length CDS of *FveRGA1* was ligated into PXY105 through Gibson cloning by using primers N-termless Bamh1BIfc_FWP & Gib_Pxy105_06210_Xba1_R. Next, full-length CDS of *FveARF8, FveARF6,* and *FveIAA4,* were amplified from cDNA and ligated into PXY106 (nYFP) at BamHI and SalI sites. BiFC constructs were transformed into *Agrobacterium* EHA105. Transformants were harvested when OD_600_ reached 1.0, and resuspended in 1/2 MS medium (suppled with 50uM acetosyringone) to a final OD_600_ of 0.5. The agrobacterial cells containing the cYFP fusion vector were mixed with agrobacterial cells containing the nYFP fusion vector at 1:1 ratio. The mixture was infiltrated into young *Nicotiana benthamiana* leave. After 48 hours, the YFP florescence signal was visualized and photographed using Leica SP5X Confocal microscope (Leica Co. USA).

For the yeast two-hybrid (Y2H), full-length CDS of *FveARF8, FveARF6, FveGID1c* were PCR amplified and ligated into GAL4 AD vector PGADT7 (Clontech Inc.) at the BamHI/XhoI, BamHI/XhoI, EcoRI/XhoI double digestion sites, respectively. Full-length CDS of *FveIAA4* and *FveGID1c* were ligated into GAL4 BD vector PBGK T7 (Clontech Inc.) through EcoRI/SalI double digestion sites. FveRGA1-C was cloned into the Y2H vector pGBTK7 by Gibson assembly using primer pair GBTK7_M5_06210_F/

GBTK7_BamH1_06210_R (Data S2). To delete the SAW domain from FveRGA1-C, primer pair RGA1-564-BglII-F & RGA1-564-SalI-R were used to amplify FveRGA1-C (-SAW) fragment and inserted into pGBK T7 at BamHI-SalI sites. The auto activity of each BD construct was examined by co-transformation with an empty AD vector PGAD T7.

To make truncated *FveARF8* for Y2H, FveARF8 (DBD), FveARF8 (MR), FveARF8 (PB1), and FveARF8 (MR+PB1) fragments were amplified respectively by primer pairs ARF8-CDS-F1/ARF8-DBD-XhoI-R; FveARF8-MD-F/ARF8-MD-R; FveARF8-PBI-F/ARF8-CDS-R1, FveARF8-MD-F/ARF8-CDS-R1 (Data S2) and inserted into PGAD T7 via EcoRI-XhoI/SalI sites. FveARF8 (DBD+MR) was amplified by ARF8-CDS-F1/ARF8-MD-R and inserted into PGAD T7 via BamHI/BglII-XhoI/SalI sites.

For Y2H essay, yeast constructs were transformed into PJ69-4A yeast strain using LiAc/PEG method (Gietz and Schiestl, 2007). The transformed yeast cells were plated onto -LW two drop-out or -HLWA four drop-out medium; -HLWA medium is often supplemented with 5mM 3-AT to reduce background. Yeast cells were grown at 30 °C for 3-4 days.

Each BiFC, Y1H, and Y2H experiment was performed three times. All repeats yielded the same result.

### Co-immunoprecipitation of proteins from yeast

The co-IP assay was performed following a published protocol (Gerace and Moazed, 2014) with minor modifications. The yeast strains containing PGBK T7-ARF6-Myc and PGAD T7-ARF8-HA (or PGBK T7-ARF6-Myc and PGAD T7-ARF6-HA) were grown to an OD_600_=1 at 28 ^0^C. After centrifugation, the yeast cells were suspended in 1×lysis buffer (100mM Na-HEPES, pH7.5, 400mM NaoAc, 2mM EDTA, 2mM EGTA, 10mM MgOAc, and 10% Glycerol) and lysed by Mini-Beadeater (model: 2412PS-12W-B30), Minebea Co., LTD. For each immunoprecipitation, 30μl Magnetic Dynabeads with protein A (Invitrogen) were resuspend in 1×lysis buffer and incubated with 5μg c-Myc primary antibody (sc-40, Santa Cruz Biotechnology) for 10min at room temperature. The protein A/c-Myc antibody slurry were dispensed into each cell lysate and rotate for 2h at 4°C. Then, the Dynabeads-cMyc antibody-protein complex were washed 5 times with wash buffer (100mM Na-HEPES, pH7.5, 400mM NaoAc, 2mM EDTA, 2mM EGTA, 10mM MgOAc, 0.25% NP-40, 3mM DTT, 1mMPMSF, and 10% Glycerol), followed by elution with 4×SDS-PAGE Protein Loading Buffer (10% SDS, 500Mm DTT, 50% Glycerol, 500mM Tris-HCL and 0.05% bromophenol blue dye) and loaded onto SDS-PAGE gel. 10% lysate was used as the “Input” loading. The western blots were performed following General V3 Western Workflow Blotting Protocol (Bio-rad, Laboratories, Inc). The anti-HA primary antibody (SKU: H6908, Sigma-Aldrich, Inc. USA) was used to detect the pull-down products in western. The co-IP experiment was repeated three times with the same result.

### Yeast one-hybrid and the luciferase reporter assays

For yeast one hybrid assay, a 1077bp promoter fragment upstream of *FveGiD1c* (gene22353) was amplified with primer FvGID1a-P-KpnI-F and FvGID1a-P-SalI-R (TableS2) and ligated into pAbAi vector (Clontech Inc.) at KpnI and SalI sites. The resulting GiD1c reporter vector was linearized by BstBI, and then introduced into competent yeast cell Y1HGold to integrate at the *ura3-52* locus. All procedures were based on the protocol Matchmaker Gold Yeast One-Hybrid Library Screening System User Manual (PT4087-1). The AD fusions in pGAD T7 with *FveARF8* (and various truncations) and *FveARF6* are the same as those described for Y2H.

The pENTR vectors LEI02 and LEI04 (Figure S9) for LUC experiments are modified from pLAH-R4R3-VP64Ter (Taylor-Teeples et al., 2015; Zhan et al., 2018). Specifically, full length VP64-35S terminator was PCR-amplified from pLAH-R4R3-VP64Ter with primers VP64-F and T35S-R (Data S2), and inserted into pCR™8/GW/TOPO™ vector (Invitrogen™) via EcoRI, to generate LEI01 vector. The LEI01 vector was digested by PacI and then self-ligated to drop off the VP64 sequence to produce LEI02. To construct the negative control vector LEI04, YFP was amplified from pLAH-L1L4-Citrine and then inserted into LEI02 vectors via EcoRI and AscI. Full length *FveARF8* or *FveARF6* was PCR amplified and inserted into LEI02 at AscI and PacI to yield *35S::FveARF8* and *35S::FveARF6* respectively. The expression cassettes of *35S::FveARF8* (or *35S::FveARF6), pGiD1c::LUC,* and *35S::REN* were simultaneously incorporated into the destination vector pLAH-LARm(Taylor-Teeples et al., 2015) through gateway cloning, resulting in the final vectors 35S::FveARF8_pGiD1c::LUC_35S::REN and 35S::FveARF6_pGiD1c::LUC_35S::REN. The 35S::Citrine_pGiD1c::LUC_35S::REN vector served as a control.

The luciferase reporter assay was performed following the instruction of Dual-Luciferase^®^Reporter Assay System (Promega, Inc, USA). Various constructs were transformed into *Agrobacterium* strain EHA105 and infiltrated into tobacco *Nicotiana benthamiana* leaves. 1μM GA was also applied during infiltration to increase the autoactivity of GiD1c. After two days, a small leaf disc from infection site was collected and finely ground in the passive lysis buffer (1×). 10 μl supernatant was mixed with LAR solution (from the Promega kit), luciferase activities were measured using TD-20/20 Luminometer (Turner Designs, Inc, USA). The LUC readout was divided by the REN readout to provide a normalized expression level. The LUC experiment was performed three times with same results. The data shown is one of the three experiments.

### RNA extraction and qRT-PCR

For testing *FveGID1c* expression (Figure 5), total RNA was extracted from YW5AF7, *arf8-1,* Hawaii 4 (H4), and *rga1-1* receptacles at stages indicated. Achenes were removed from the receptacle, and 3 to 5 receptacles from several plants of the same genotype were combined as one biological replicate. The CTAB RNA extraction method was used(Gambino et al., 2008). The CTAB RNA extraction solution (2% CTAB, 2% PVPk-30, 10mM Tris-Hcl(pH8), 25mM EDTA, 2M Nacl, 0.5g/L Spermidine) was added to the flesh tissues. The tissues were ground immediately following by the addition of 2/3 volume of 10M LiCl to precipitation RNA. In total, 1 ug total RNA was used for cDNA synthesis with SuperScript IV VILO Master Mix (Invitrogen, USA). The cDNA product was diluted 4 times and ~5 ul was used as a template in qRT-PCR using BioRad CFX96 Real-time system and SYBR Green PCR MasterMix (Applied Biosystems). The expression levels of four *F.vesca* genes FveARF8, FveARF6, FveIAA4, and FveGiD1c were quantified via 4 primer pairs ARF8-F1-2/ARF8-R1-2, FveARF6-F1/FveARF6-R1, FveIAA4-F1 /FveIAA4-XhoI-R and Fv GID1c-F2/ Fv GID1c-R2 (Data S2), respectively. *F. vesca* gene03773 was used as the internal control with primers gene03773-F/gene03773-R (Caruana et al., 2018). The experiment was performed three times each with RNA extracted from newly collected samples (ie. biological replicates).

### Mining consensus co-expression network and other bioinformatics tools

The expression pattern of each *F. vesca* gene in wild type (YW5AF7) was visualized via strawberry eFP browser (Hawkins et al., 2017) (http://mb3.towson.edu/efp/cgi-bin/efpWeb.cgi). The consensus co-expression network analysis of YW5AF7 floral and fruit tissues(Shahan et al., 2018) was mined using the network website (http://159.203.72.198:3838/fvesca/). The co-expression cluster/module was visualized using Cytoscape (cytoscape.org). Protein alignments were performed using clustalw (https://www.genome.jp/tools-bin/clustalw). Heatmap was generated with morpheus (https://software.broadinstitute.org/morpheus/).

### Tissue section and histological staining

Tissue section was performed following the protocol (Retamales and Scharaschkin, 2014) with minor modification. YW5AF7, *arf8-1, rga1-1* fruit (receptacle and achene) of each stage was fixed in FAA (50% ethanol, 5% acetic acid, 3.7% Formaldehyde) for 3 days and went through an ethanol series 50%, 60%, 70%, 80%, 90%, 95% (containing 0.1% Eosin), and 100%. The tissues went through a series of changes in 25% xylenes/75% ethanol, 50% xylenes/50% ethanol, 75% xylenes/25% ethanol, and finally 100% xylenes. After two more changes of 100% xylene, paraffin chips were added into the xylene. Afterwards, the tissue in xylene/paraffin mix is exchanged with 100% paraffin wax. After 4 changes of 100% paraffin in a duration of 4-5 days, a wax boat was poured. The tissues were sectioned at 12 μm thickness on a microtome, and stained by Safranin-O/Fast Green based on a protocol (http://microscopy.berkeley.edu/Resources/instruction/staining.htm). Slides were mounted with Permount (Fisher) and dried overnight.

## Acknowledgements

This work has been supported by a grant from the National Science Foundation (IOS 1444987). We thank Dr. Ramin Yadegari at University of Arizona, Samuel Hazen at UMass Amherst, Dr. Yanhai Yin from Iowa State University, and Dr. Kunsong Chen from Zhejiang University for vectors.

## Author Contributions

J. Z. and Z. L. designed experiments. J. Z., J. S., and L. G. performed experiments. X. H. and A. P. assisted experiments. J. Z. and Z. L. analyzed and discussed the data, Z. L. and J. Z. wrote the paper.

